# Immunomodulatory effects of cytokine-induced expansion of cytotoxic lymphocytes in a mouse model of lupus-like disease

**DOI:** 10.1101/815399

**Authors:** Seth D. Reighard, Durga Krishnamurthy, Hilal Cevik, David E. Ochayon, Ayad Ali, Harsha Seelamneni, Hermine I. Brunner, Stephen N. Waggoner

**Affiliations:** Center for Autoimmune Genomics and Etiology, Cincinnati Children’s Hospital Medical Center, Cincinnati, OH, USA; Medical Scientist Training Program, University of Cincinnati College of Medicine, Cincinnati, OH, USA; Immunology Graduate Program, University of Cincinnati College of Medicine, Cincinnati, OH, USA; Molecular and Developmental Biology Graduate Program, University of Cincinnati College of Medicine, Cincinnati, OH, USA; Department of Pediatrics, University of Cincinnati College of Medicine, Cincinnati, OH, USA; Division of Rheumatology, Cincinnati Children’s Hospital Medical Center, Cincinnati, OH, USA

**Keywords:** Lupus, SLE, CD8 T, NK cell, IL-15, IL-2, Immunotherapy, T_FH_

## Abstract

Certain therapies (i.e. daclizumab) that promote expansion of natural killer (NK) cells are associated with clinical amelioration of disease in the context of multiple sclerosis and associated mouse models. The clinical benefits are putatively attributable to an enhanced capacity of NK cells to kill activated pathogenic T cells. Whether a parallel approach will also be effective in systemic lupus erythematosus (lupus), a multi-organ autoimmune disease driven by aberrant responses of self-reactive T and B cells, is unclear. In the present study we assess the therapeutic impact of IL-2 and IL-15-based strategies for expanding NK cells on measures of lupus-like disease in a mouse model. Unexpectedly, cytokine-mediated expansion of cytotoxic lymphocytes aggravated immunological measures of lupus-like disease. Depletion studies reveal that the negative effects of these cytokine-based regimens can largely be attributed to expansion of CD8 T cells rather than NK cells. These results provoke caution in the use of cytokine-based therapeutics to treat co-morbid cancers in patients with lupus and highlight the need for new methods to selectively expand NK cells in order to further assess their clinical value in autoimmune disease.

## Introduction

The Centers for Disease Control and Prevention estimates that 161,000 Americans, most of whom are ethnic minorities and women of child-bearing age, suffer the debilitating consequences of systemic lupus erythematosus (or lupus). Lupus is an autoimmune disease characterized by the rampant production of autoantibodies that drive widespread organ damage^1^. As a result, lupus is the 5^th^ leading cause of death for African American and Hispanic women between 15 and 24 years of age^2^. There is no cure for lupus, and therapy is limited mostly to immunosuppressive drugs. Amongst numerous side effects, these treatments broadly dampen immune responses, causing impaired wound healing and an increased susceptibility to dangerous infections. There is a growing focus on development of immunomodulatory therapies that more selectively constrain pathogenic autoreactive lymphocytes in patients with lupus, thereby obviating the need for potentially harmful non-selective suppression of the immune system.

Multiple lines of evidence in patients with lupus and in mouse models of lupus-like disease point to dysregulation of follicular helper T cell (T_FH_) and germinal center (GC) B-cell responses as a key driver of disease pathogenesis^3-5^. The resulting aberrant GC responses facilitate development of autoantibody-secreting plasmablasts that contribute to harmful inflammation and pathogenic deposition of immune complexes in affected organs, including the kidney^1^. While clinical strategies to deplete T_FH_ are presently lacking, broad depletion of B-cells^6^ or neutralization of the B-cell activating factor^7^ show promise for ameliorating lupus disease. These results suggest that other immunotherapeutic strategies that target T_FH_ and B cells may be able to effectively alleviate lupus disease.

Our previous work revealed that natural killer (NK) cells exhibit potent immunosuppressive capacity that limits T_FH_ and GC responses resulting from infection or vaccination^8,9^. This activity involves perforin-dependent lysis of a fraction of activated CD4 T cells by NK cells, resulting in reduced antiviral CD8 T cell and humoral immune responses^8-10^. Unfortunately, the frequency and function of circulating NK cells is markedly reduced in patients with lupus^11-15^, particularly during disease flares^16,17^. Likewise, NK-cell frequencies are reduced in mouse models of lupus-like disease, including Fas-mutant *lpr* mice^18^. Of note, antibody-mediated depletion of NK cells in these mice accelerated disease, while adoptive transfer of NK cells delayed disease onset^18^. These data suggest that an inherent or acquired NK-cell numeric and functional deficiency may contribute to the pathogenesis of lupus. Moreover, the reduced NK-cell pool in the context of lupus coupled with the potent immunoregulatory function of these cells raises the specter of therapeutically expanding NK cells to alleviate lupus disease.

Several methods currently exist to semi-selectively expand endogenous NK cell populations *in vivo*^19-22^. These reagents are often based on the NK-cell stimulating activities of the cytokines interleukin-2 (IL-2) and IL-15^23,24^. Of note, both IL-2 and IL-15 engage NK cells via IL-2 beta receptor (IL-2Rβ, or CD122), which is also expressed on activated T and NKT cells^25^. Therefore, targeting of IL-2 to IL-2Rβ via daclizumab (anti-IL-2Rα antibody) or IL-2/anti-IL-2 (clone S4B6) antibody complexes (IL-2C), results in marked expansion of the NK-cell and CD8 T-cell compartments in mice and humans^23,26^. Daclizumab-mediated expansion of immunoregulatory NK cells in multiple sclerosis patients is linked to clinical lessening of disease^26,27^, whereas IL-2C treatment reduced disease measures in the MRL/*lpr* mouse model of lupus^28^. Likewise, lupus-like disease in the B6 × DBA/2 F1 mouse model was alleviated by therapeutic administration of IL-2C^29^. In contrast, pre-treatment with IL-2C prior to disease initiation resulted in worse disease. The potential contributions of both NK cells and CD8 T cells in these disparate models complicates interpretation of any potential clinical benefits of IL-2C in human lupus. Moreover, distinct functional consequences of IL-15-versus IL-2-mediated expansion of NK cells in infection models^30^ suggest that IL-15 superagonist compounds like N-803 (previously known as ALT-803^31^) may provoke greater immunoregulatory functions in expanded NK cells.

In the present study, we compare the prophylactic and therapeutic effects of both IL-2C- and N-803-induced expansion of NK and CD8 T cells in a mouse model of lupus-like disease. In this chronic graft-versus-host disease (cGvHD) model^32,33^, CD4 T cells from B6I-*H2*-*Ab1*^*bm12*^/KhEgJ (bm12) donor mice undergo alloactivation following transfer into C57BL/6 mice due to three amino acid mismatches between their major histocompatibility complex (MHC) class II I-A^b^ molecules. The resulting cGvHD provokes numerous immunological manifestations resembling lupus, including robust T_FH_ and GC B-cell responses, elaboration of antinuclear antibodies (ANA), and development of glomerulonephritis. Surprisingly, both IL-2C and N-803 exacerbated immunological measures of disease in this mouse model. CD8 T cells, but not NK cells, were linked to immune enhancement in this disease model following cytokine treatment.

## Materials & Methods

### Mice

C57BL/6 and CD45.1^+^ (B6.SJL-*Ptprc*^*a*^*pec*^*b*^/BoyJ) breeder pairs and experimental recipient mice were obtained from The Jackson Lab. CD45.1^+^ B6(C)-*H2*-*Ab1*^*bm12*^/KhEgJ (bm12) mice were a kind gift from Dr. Edith Janssen. Male and female mice between eight and twenty weeks of age were used in experiments. All mice were maintained under specific pathogen-free conditions in accordance with guidelines from the Association for Assessment and Accreditation of Laboratory Animal Care International. Experiments were performed in accordance with ethical guidelines approved by the Institutional Animal Use and Care Committee of Cincinnati Children’s Hospital Medical Center.

### Bm12 cell transfer and tissue harvest

Spleens were harvested from B6(C)-*H2*-*Ab1*^*bm12*^/KhEgJ (bm12) donor mice and passed through 70 μm nylon strainers to obtain single-cell suspensions. Following enumeration using 3% acetic acid with methylene blue (StemCell Technologies), splenocytes were resuspended in phosphate-buffered saline and forty to sixty million cells were injected either intraperitoneally or intravenously (via retro-orbital injection under isoflurane anesthesia) into each recipient C57BL/6 mouse. This range of donor cells falls above the necessary minimum dose of thirty million splenocytes previously determined by our colleagues and elsewhere noted^32^. Although early experiments utilized an intraperitoneal graft injection that is traditional to the disease model, later experiments used intravenous delivery due to an observed enhanced consistency of engraftment that did not significantly affect downstream disease measures (data not shown). Donor and recipient mice were age-matched. Female-derived graft was injected into female or male hosts, or male-derived graft into male hosts (but not male-derived grafts into female hosts so as to avoid Y chromosome-mediated rejection). When necessary, whole blood was obtained via laceration of the tail vein and collected into either EDTA or SST Microtainer tubes (BD Biosciences). Prior to staining for flow cytometry, whole blood from EDTA tubes was subjected to red blood cells lysis via ACK Lysing Buffer (Thermo Fisher). Following euthanasia, mouse spleens were weighed and processed into single-cell suspensions (as above), splenic red blood cells were lysed (as above), and remaining splenocytes were enumerated using either Trypan Blue exclusion or 3% acetic acid with methylene blue prior to staining for flow cytometry. Whole blood was collected through the inferior vena cava and serum was isolated using SST Microtainer tubes. Due to unknown factors in our vivarium, immune perturbation and ANA antibody production did not lead to measurable kidney damage or proteinuria in any of the mice assayed in this manuscript.

### Targeted cytokine treatment

For each mouse, 1 μg of recombinant murine IL-2 (PeproTech) was combined with 10 μg anti-mouse IL-2 (clone S4B6; BioXCell) in 200 μL Hank’s Balanced Salt Solution (HBSS), then incubated overnight at 4°C while mixing. For each control mouse, 1 μg of recombinant murine IL-2 was incubated overnight with 10 μg rat IgG2a (BioXCell) in 200 μL HBSS. As a pre-disease treatment, each mixture was intraperitoneally injected every other day for a total of 3 injections, after which a bm12 splenocyte graft was injected the following day (see above). As a post-disease treatment, injections were administered once daily for three days beginning one month after graft injection^29^.

Experimental mice received 200 μg/kg of N-803 (provided via collaboration with Altor Biosciences, now part of NantWorks LLC) in PBS injected subcutaneously, while control mice received a molar equivalent of recombinant human IL-15 (Peprotech) in PBS. As a pre-disease treatment, two injections were administered, once every other day, finishing one day prior to bm12 graft injection. As a post-disease treatment, injections were administered once weekly over the course of 3 months, beginning 1 week after bm12 graft injection (for a total of 11 treatments).

### Administration of cell-depleting antibodies

For NK cell, CD8 T cell, or mock depletion, mice were intraperitoneally injected once with either 25 μg *InVivo*MAb anti-mouse NK1.1 (BioXCell, clone PK136)^10^, 100 μg *InVivo*MAb anti-mouse CD8α (BioXCell, clone YTS169.4), or 25 μg *InVivo*MAb mouse IgG2a isotype control (BioXCell, clone BE0085) in HBSS, respectively. Antibodies were administered one day prior to the first treatment with either IL-2/αIL-2 complexes, N-803, or their respective controls.

### ELISA assays

Mouse serum or plasma was assessed via the IgG (Total) Mouse ELISA Kit (Thermo Fisher), according to the manufacturer’s directions, using 96-well Nunc Maxisorp plates (Thermo Fisher) with samples diluted 1:20,000 – 1:40,000. Serum or plasma was assessed for ANA using the Mouse ANA Total Ig ELISA Kit (Alpha Diagnostic International), according to the manufacturer’s directions, with samples diluted 1:50. Total IgG and ANA ELISA plates were read using a GlowMax Discover Microplate Reader (Promega) on the absorbance setting at 450nm.

### Flow cytometry

Fluorescently-conjugated antibodies used were obtained from either Biolegend, Ebioscience (ThermoFisher), Invitrogen, or BD Biosciences. For surface staining, cells were resuspended at a concentration of 1-2×10^6^/mL in 50-100 µL of cold HBSS containing 5% heat-inactivated FBS (Gibco), 100U/mL penicillin, and 100 µg/mL streptomycin (“FACS buffer”). Fluorescently-conjugated antibodies were each added at the manufacturer’s recommended concentration. Cells were incubated at 4°C for 20 minutes, then washed twice with FACS buffer, and either analyzed fresh (same-day) or fixed using 100 µL Cytofix (BD Biosciences) for 20 minutes at 4°C. All fixed cells were washed twice with FACS buffer to remove fixative and kept at 4°C in FACS buffer until analysis (1-3 days later). Acquisition of stained cells was performed using a Fortessa or LSRII cytometer (BD Biosciences) with FACSDiva software (BD Biosciences).

### Statistical analyses

All flow cytometric analysis was performed using FlowJo v.10 software. Compensation was performed using single-color-stained cells and/or UltraComp eBeads (ThermoFisher), while Zombie dye single-color controls were made using cells pre-heated on a heat block at 70°C for 10 minutes. All electronic gating was performed downstream of a forward scatter area by forward scatter height “singlet” gate. ELISA standard curves were generated using the linear regression function in GraphPad Prism, during which analyte concentrations were interpolated based on OD values, then calculated based on dilution factor.

All statistical analysis was performed using GraphPad Prism 8. Normal distribution of data was assessed by Kolmogorov-Smirnov test. Significant differences between two datasets exhibiting normal distribution were assessed by unpaired, two-tailed Student’s T-test, while non-normally distributed datasets were compared using Mann Whitney U test. Comparisons among multiple groups made via Kruskal-Wallis test with Dunn’s multiple comparison testing of group means. Nonparametric Spearman correlation was used to assess relationships between independent measurements.

## Results

### The bm12 cGvHD model is associated with contraction of the NK-cell compartment

Injection of splenocytes from CD45.1^+^ bm12 mice (graft) into CD45.2^+^ C57BL/6 recipient (host) mice results in marked splenomegaly characterized by activation and expansion of graft PD1^+^ CXCR5^+^ CD4^+^ T_FH_ cells, host Fas^+^ GL7^+^ CD19^+^ GC B cells, and host CD138^+^ CD19^+^ plasmablasts relative to C57BL/6 receiving a control graft of BoyJ splenocytes (**Figure 1A**). Despite the increase in splenocyte count during this cGvHD response, the numbers of host CD3^-^ NK1.1^+^ NKp46^+^ NK cells were 60% reduced (p=0.0002) in the spleen (**Figure 1B**). This contraction of the NK-cell compartment is consistent with that observed in other mouse models of lupus-like disease ^18^ as well as in patients with lupus^11-15^. In addition to quantitative reductions, host CD3^-^ NK1.1^+^ NKp46^+^ splenic NK cells from bm12-engrafted mice exhibited higher surface expression of CD27 (*Tnfrsf7*), NKG2A (*Klrc1*), Tumor necrosis factor-related apoptosis-inducing ligand (TRAIL/*Tnfsf10*), and CD11c (*Itgax*) than mice injected with a CD45.1^+^ control graft (**Figure 1C**). This phenotype (CD27^+^, NKG2A^+^, TRAIL^+^) is consistent with more immature NK cells^34^, while the increased expression of CD11c is characteristic of an atypical, pro-inflammatory NK-cell subset observed in patients with lupus and certain mouse models of lupus-like disease^35-37^.

**Figure 1.**
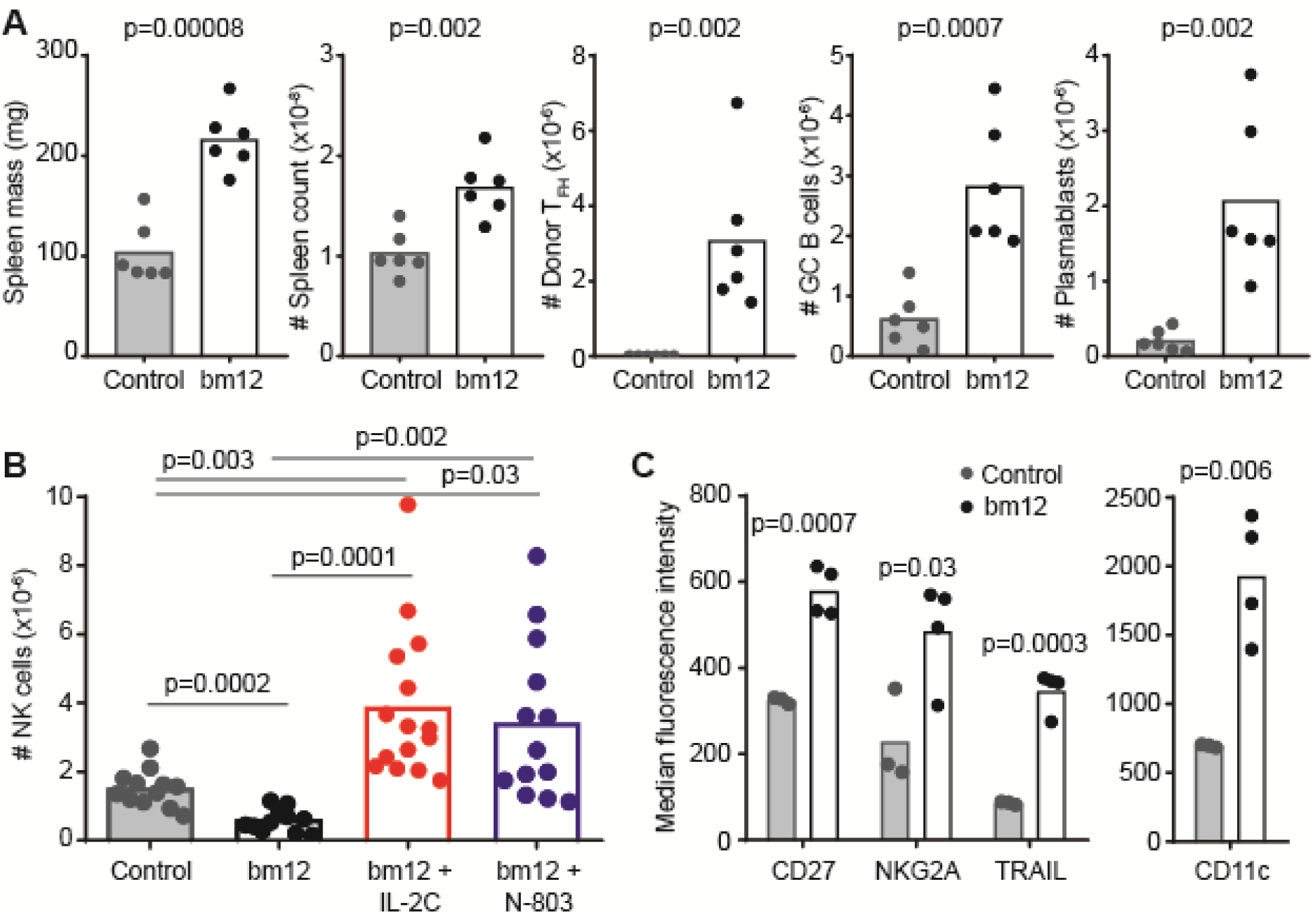
Cytokine-mediated reversal of NK-cell attrition associated with bm12 lupus model. C57BL/6 mice received a graft of CD45.1^+^ (Control) or CD45.1^+^ bm12 splenocytes and were evaluated two weeks later. (**A**) Spleen masses, spleen cellularities, donor CD45.1^+^ CXCR5^+^ PD-1^+^ CD4^+^ T_FH_ counts, host CD19^+^ Fas^+^ GL7^+^ GC B cell counts, and host CD19^+^ CD138^+^ plasmablast counts are displayed (n=6 mice/group). (**B)** Total splenic NK cells (CD3^-^ NK1.1^+^ NKp46^+^) in control- and bm12-engrafted mice (n=12-15/group) that were pre-treated with either IL-2C or N-803 as described in methods. (**C**) Cell surface phenotype of NK cells in control and bm12 recipient mice (n=3-4/group) determining by flow cytometric assessment of receptor staining intensity. Results are representative of two or more experimental replicates.

### Disease-associated attrition of NK cells can be reversed with targeted cytokine treatment

Injections of IL-2C (IL-2 complexed with S4B6 anti-IL-2 antibody) or the IL-15 superagonist N-803 into C57BL/6 mice results in expansion of the NK-cell and CD8 T-cell compartments, while Foxp3^+^ CD4^+^ T regulatory cell proportions remained relatively unchanged^38,39^. To determine if NK-cell attrition in the bm12 model can be averted via cytokine-mediated expansion of the NK-cell compartment, we treated C57BL/6 mice with IL-2C three times over a span of six days prior to initiation of disease model. Pre-treatment with IL-2C before injection of bm12 cells resulted in a >6-fold increase in host splenic NK-cell frequencies relative to control-treated (IL-2 and rat IgG2a) mice receiving a bm12 graft (**Figure 1B**). Likewise, pre-treatment with N-803 promoted a >5-fold increase in the frequency of host splenic NK cells relative to untreated bm12-engrafted mice. Of note, NK-cell frequencies in IL-2C- or N-803-treated mice were >2-fold greater than in control mice given a non-disease inducing graft of CD45.1^+^ splenocytes (**Figure 1B**). N-803 pretreatment further increased up-regulation of CD27 (IL-15: 1945±404 MFI, N-803: 3401±656 MFI, n=4/group, p=0.004) and NKG2A (IL-15 218±44 MFI, N-803: 1120±88 MFI, n=4/group, p=0.002) on NK cells after bm12 cell transfer, but had no added effect on expression of TRAIL or CD11c (data not shown). Similar observations were made with IL-2C (data not shown). Thus, cytokine-mediated expansion of NK cells prior to bm12 cell transfer can prevent or even reverse model-associated attrition of NK cells.

### IL-2C pretreatment transiently exacerbates cGvHD

The diminished number of potentially immunoregulatory NK cells^8-10,26,40-51^ in the bm12 model of disease (**Figure 1B**) may contribute to dysregulated responses of T and B cells leading to autoimmune disease. Accordingly, the marked expansion and maintenance of high number of NK cells after IL-2C pre-treatment (**Figure 1B+2A**) should result in diminished measures of cGvHD and lupus-like disease. In contrast to expectations, the number of donor T_FH_, host GC B cells and plasmablasts in the spleen of IL-2C-treated, bm12-engrafted C57BL/6 mice two weeks after disease initiation was elevated relative to mice treated with IL-2 and non-complexed irrelevant IgG2a (**Figure 2A**). CD8 T cell counts were also elevated at the two week time point (Control: 6.4±1.9×10^6^ CD8 T-cell, IL-2C: 10.4±2.6×10^6^, n=6-7, p=0.01). Total serum immunoglobulin (IgG) levels were elevated relative to baseline (as expected at this time point), but to similar levels in IL-2C-treated and control bm12-recipient mice (**Figure 2A**). Neither the expansion of NK cells and host CD45.2+ CD8 T-cells (Control: 15.1±2.2×10^6^, IL-2C: 22.2±11.1×10^6^, n=6-7, p=0.1) nor the enhanced immunological measures of disease were not sustained at three month time point, when IL-2/S4B6 pre-treated and control-treated mice exhibited similar numbers of donor T_FH_, host GC B cells and plasmablasts (**Figure 2B**). Comparable titers of total IgG (not shown) and antinuclear antibodies (ANA), which become measurable at this time point, were also observed in sera of IL-2/S4B6 pre-treated mice relative to controls (**Figure 2B**). Thus, increased immunological cGvHD measurements at the two-week time point in IL-2/S4B6 pretreated mice (**Figure 2A**) were not associated with measurable amplification of later measures (**Figure 2B**) of immune disease in this model.

**Figure 2.**
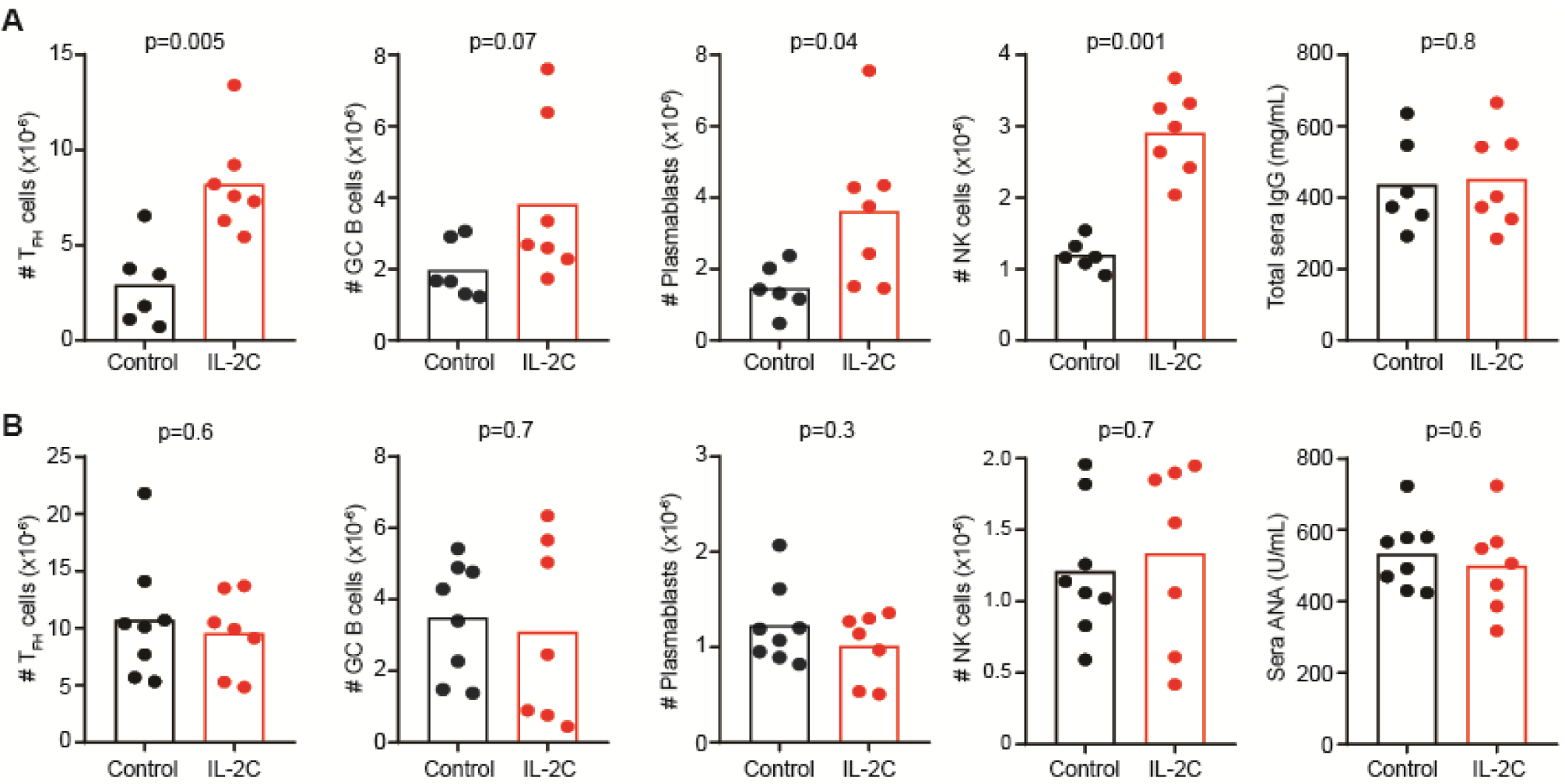
IL-2C pretreatment transiently exacerbates cGvHD. C57BL/6 mice were given three injections of IL-2C (IL-2/S4B6) or IL-2/IgG2a (Control) over span of 6 days prior to infusion of bm12 splenocytes. The number of donor CD45.1^+^ CXCR5^+^ PD-1^+^ CD4^+^ T_FH_, host CD19^+^ Fas^+^ GL7^+^ GC B cells, CD19^+^ CD138^+^ plasmablasts, and NK1.1^+^ NKp46^+^ CD3^-^ NK cells in recipient spleens (n=7-8/group) was evaluated by flow cytometry (**A**) two weeks or (**B**) three months after disease initiation. Sera IgG and ANA titers evaluated by ELISA at two week and three month time points, respectively. Results are representative of at least two experimental replicates performed for each measurement.

### Therapeutic application of IL-2C exacerbates lupus-like disease in mice

To assess whether therapeutic cytokine-mediated expansion of NK cells can alleviate established lupus-like disease, control and bm12-recipient mice were administered daily therapeutic regimens of IL-2C for 3 consecutive days starting one month after disease initiation, an approach that parallels a prior study using this complex in a similar disease model^29^. Two-months after therapy initiation, IL-2C-treated mice exhibited increased host splenic plasmablast counts and elevated sera ANA titers relative to control treated mice (**Figure 3**). Non-significant trends for elevations in host GC B-cell counts and total sera IgG expression levels were also observed. Thus, similar to pre-treatment, therapeutic application of IL-2C exacerbates measures of disease in this model.

**Figure 3.**
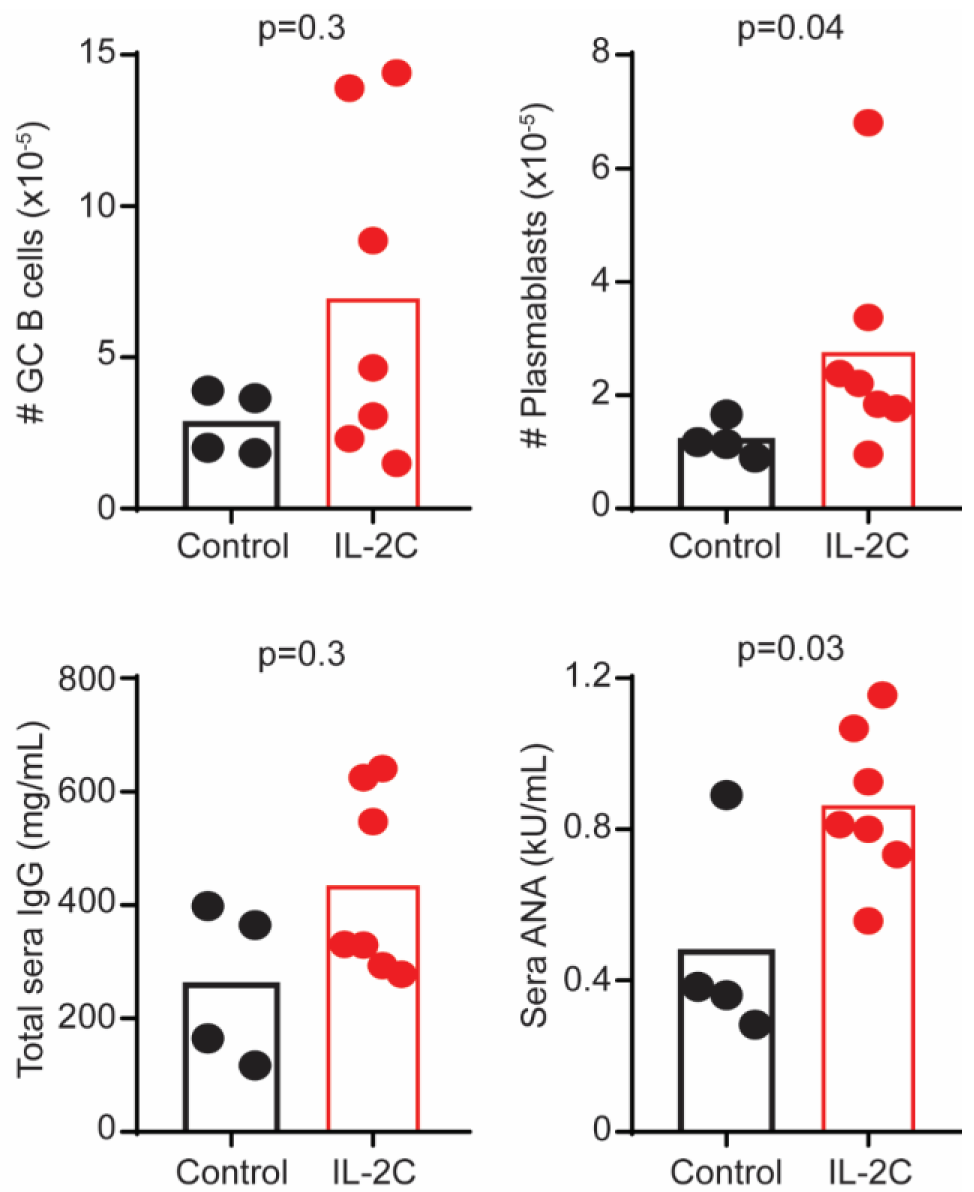
Therapeutic application of IL-2C aggravates bm12-induced disease. C57BL/6 mice were administered bm12 splenocytes one month prior to application of three daily injections of IL-2C (IL-2/S4B6) or IL-2/IgG2a (Control). The numbers of host splenic CD19^+^ Fas^+^ GL7^+^ GC B cells and CD19^+^ CD138^+^ plasmablasts as well as sera titers of IgG and ANA were evaluated (n=4-7 mice/group) two months after IL-2C therapy (third month of disease). Results are representative of at least two experimental replicates.

### N-803 pretreatment transiently exacerbates cGvHD

Since IL-15-based expansion can more robustly trigger accumulation of NK cells with greater immunoregulatory capacity than is seen with IL-2C-mediated expansion ^30^, especially when using the novel IL-15 superagonist N-803 ^52^, we tested the hypothesis that N-803 pre-treatment would reduce disease measures in the bm12 model. Unfortunately, donor T_FH_, host GC B cell, CD8 T cells, and plasmablast counts were also elevated in diseased mice pretreated with N-803 relative to control mice treated with molar-equivalent IL-15 (**Figure 4A**). There was also a non-statistically significant trend towards increased sera IgG titers at two weeks in N-803 pre-treated mice (**Figure 4A**). The measured increases in T_FH_, GC B cell, plasmablasts, and CD8 T cells were short-lived, as analysis 3-months post-disease initiation revealed similar numbers of these cells and comparable ANA titers in N-803 pre-treated mice relative to control mice (**Figure 4B**). Elevated NK-cell frequencies with N-803 treatment at 2 weeks (**Figure 1B**) also returned to baseline (IL-15: 3.1±2.3×10^6^, N-803: 2.6±1.5×10^6^, n=8-9, p=0.7).

**Figure 4.**
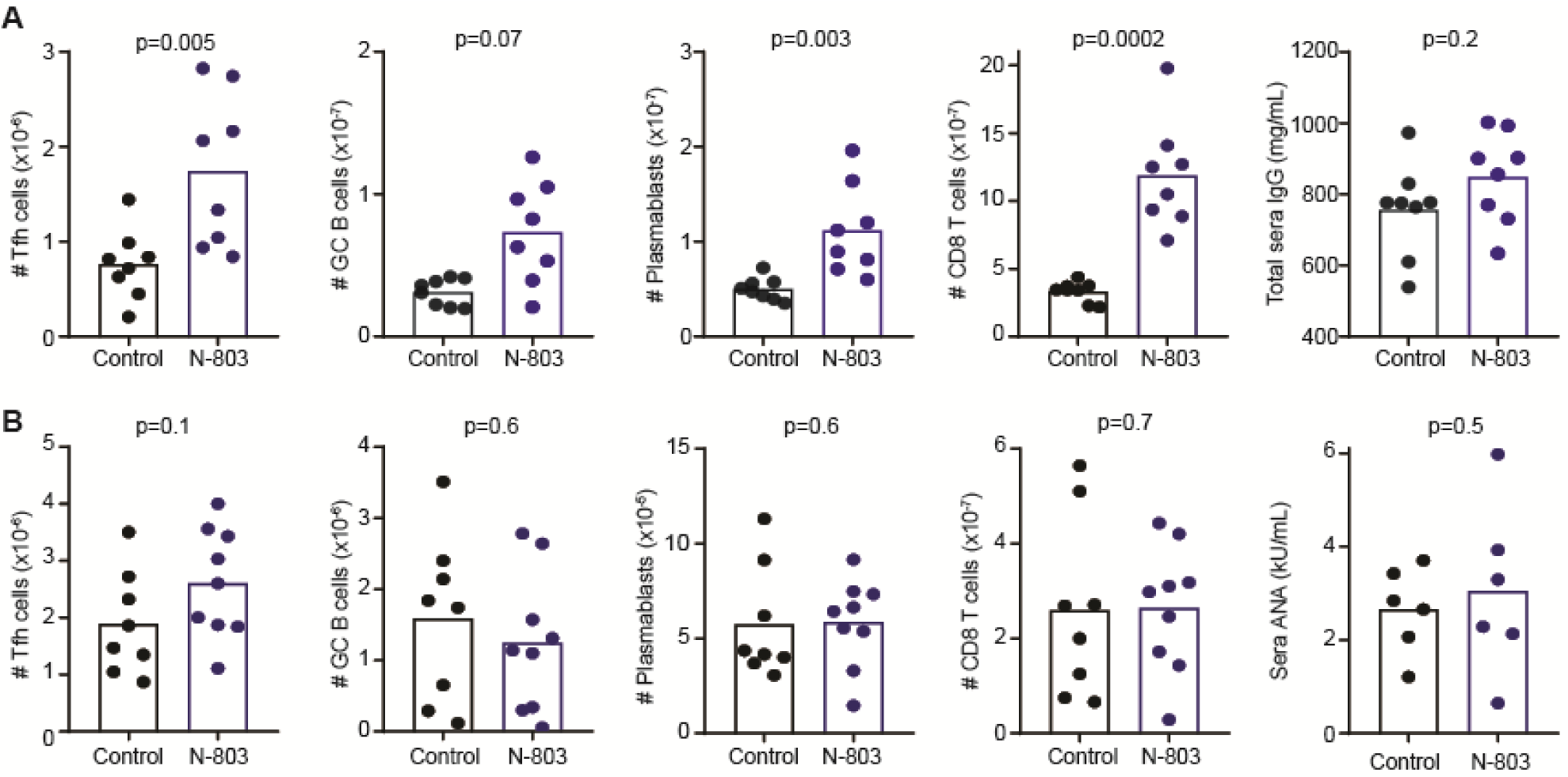
N-803 pretreatment transiently exacerbates bm12-induced disease. C57BL/6 mice were given two injections of 200 μg/kg N-803 or IL-15 (Control) over span of 4 days prior to infusion of bm12 lymphocytes. Splenic CD45.1^+^ CXCR5^+^ PD-1^+^ CD4^+^ T_FH_, host CD19^+^ Fas^+^ GL7^+^ GC B cells, CD19^+^ CD138^+^ plasmablasts, and host CD45.2^+^ CD3^+^ CD8^+^ T cells as well as sera IgG and ANA titers were assessed (**A**) two weeks (n=8/group) or (**B**) three months (n=8-9/group) after disease initiation. ANA was only assessed in mice from whom sufficient quantity of blood was obtained. Similar results were obtained in two independent experimental replicates in A, while replicates were pooled in B.

### Therapeutic application of N-803 exacerbates immune responses in this lupus model

To assess whether N-803 treatment can alleviate established lupus-like disease, a clinically relevant regimen ^53,54^ of weekly N-803 or control IL-15 injections were initiated one week after bm12 splenocyte infusion and continued for up to eleven weeks. Early (1 to 2 months) after therapy initiation, there was a trend for increased donor (CD45.1_+_) T_FH_ cell as well as host, plasmablast, GC B cell, and CD8 T cell frequencies in mice treated with N-803 therapeutically relative to control-treated mice (**Figure 5A**). Due to pandemic interruptions, replicate experiments spanned from 1 to 5 months post-treatment initiation. Collective analysis of treated and control mice reveals that the numbers of splenic CD8 T cell across this timespan shows a positive correlation with both the number of splenic plasmablasts and donor T_FH_ (**Figure 5B**). By contrast, NK-cell frequencies exhibit a weak negative correlation with donor T_FH_ frequencies (**Figure 5C**). Therapeutic N-803 treatment was also associated with non-significant (p>0.1) increases in sera ANA titers 1-2 months (Control: 4.5±1.8 kU/mL; N-803: 6.4±2.8 kU/mL, n=6-7) and 3-5 months (Control: 4.4±1.6 kU/mL; N-803: 5.6±2.5 kU/mL, n=10-11) after treatment initiation. Collectively, these data suggest N-803 therapy after disease initiation mildly exacerbates immune measures of disease in this model, and that this phenomenon may associate with an expanded CD8 T-cell compartment.

**Figure 5.**
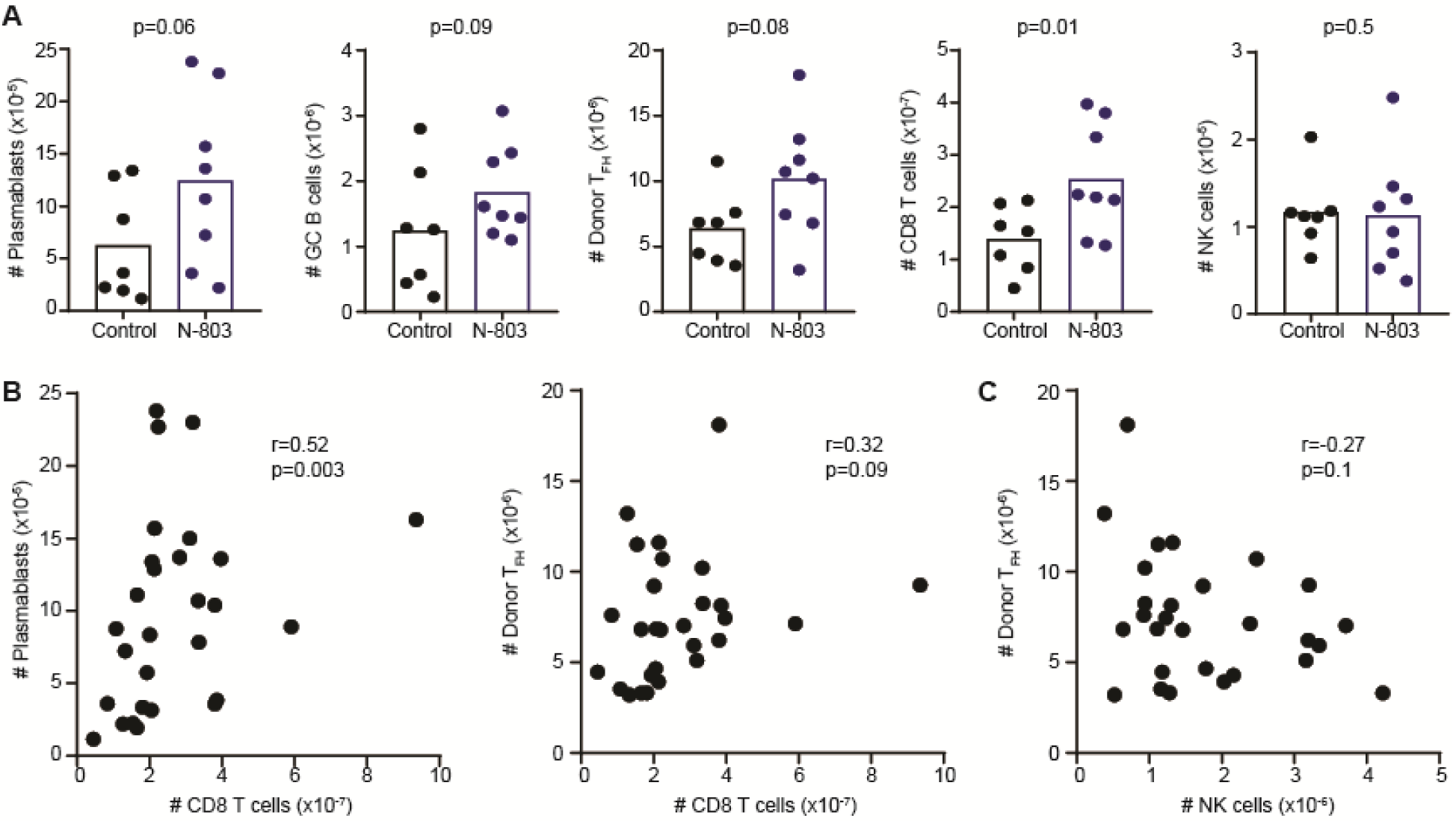
N-803 therapy augments bm12-induced disease measures. C57BL/6 mice were administered bm12 splenocytes one week prior to application of onset of weekly injections of 200 μg/kg N-803 or IL-15 (Control). (**A**) The number of donor CD45.1^+^ CXCR5^+^ PD-1^+^ CD4^+^ T_FH_ cells and CD19^+^ Fas^+^ GL7^+^ GC B cells, CD19^+^ CD138^+^ plasmablasts, host CD45.2^+^ CD3^+^ CD8^+^ T cells, and NKp46+ NK1.1 CD3- NK cells in spleen was assessed in groups of mice (n=7-8/group) at one to two months post-disease initiation. Spearman correlation analysis in mice assessed across 5 month timespan showing association between (**B**) CD8 T cells and plasmablast or T_FH_ frequency as well as between (C) NK cell and T_FH_ frequencies. Results are pooled from replicate experiments.

### CD8 T cells, but not NK cells, contribute to N-803-mediated enhanced B-cell responses

Cytokine pretreatment-induced expansion of CD8 T cells and NK cells temporally associated with enhanced immunological measures of disease in this mouse model (**Figure 2&4**), yet only CD8 T cell frequencies showed positive correlations with these parameters with therapeutic cytokine expansion (**Figure 5**). Therefore, we sought to evaluate the dueling contribution of NK cells and CD8 T cells to enhanced disease measures by depleting each of these cell types with anti-NK1.1 or anti-CD8α antibodies, respectively. Thereafter, depleted and non-depleted mice were pre-treated with N-803 for one week prior to engraftment of bm12 splenocytes. Both N-803 and IL-2C-stimulated expansions of CD8 T or NK cells were ablated by depletion of CD8 T cells (**Figure 6A, D**) or NK cells (**Figure 6B, E**), respectively, without affecting the opposing population of cells. Importantly, host GC B-cell responses were increased following N-803 pre-treatment of bm12-engrafted mice in non-depleted control mice (p=0.002) and in mice depleted of NK cells (p=0.003) prior to N-803 administration (**Figure 6C).** However, this increase was ameliorated (p=1.0) by CD8 T-cell depletion prior to N-803 therapy. In a similar fashion, CD8 T-cell but not NK-cell depletion resulted in normalization of GC B-cell responses after IL-2C treatment (**Figure 6F**). Thus, cytokine-induced increases in CD8 T-cell frequencies in the bm12 lupus-like cGvHD model are associated with enhanced immunological measures of disease.

**Figure 6.**
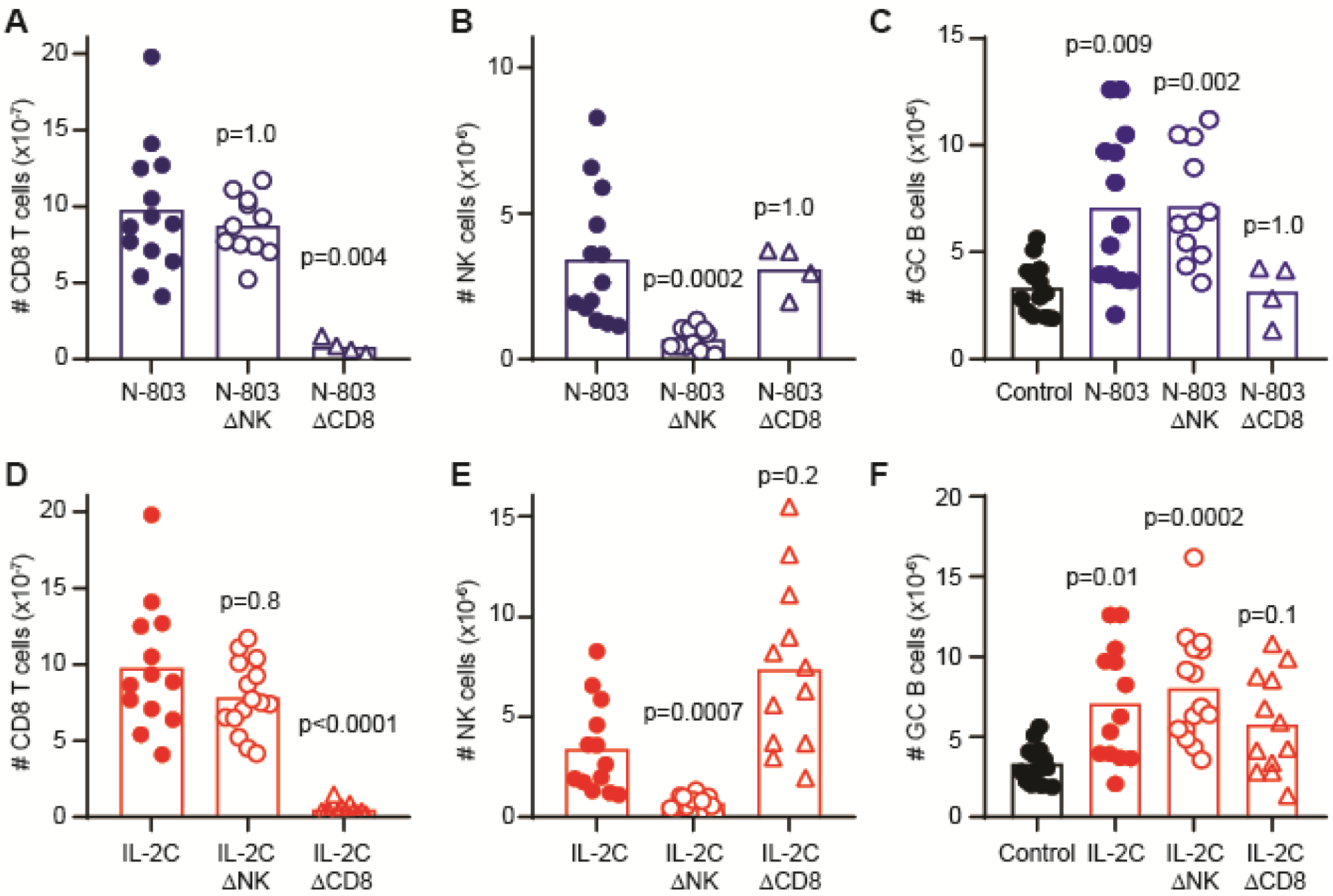
Contribution of CD8 T cells to cytokine augmentation of immune disease. C57BL/6 mice (n=4-13/group) were treated with IgG2a isotype control, 25 μg anti-NK1.1 antibody (ΔNK), or 100 μg anti-CD8α antibody (ΔCD8) one day prior to (**A-C**) N-803, (**D-F**) IL-2C, or appropriate Control treatments as described in methods. (**A**,**D**) CD3^+^ CD8^+^ T cell, (**B**,**E**) CD3^-^ NK1.1^+^ NKp46^+^ NK cell, or (**C**,**F**) CD19^+^ Fas^+^ GL7^+^ GC B cells counts in spleen was determined by flow cytometry two weeks after bm12 disease initiation in groups of mice (n=4-13/group). Statistically significant changes over time points or between treatment groups determined by Kruskal-Wallis test with Dunn’s multiple comparison testing. Results are pooled from two to three independent experiments.

## Discussion

The investigation herein sought to determine whether IL-2C or N-803-mediated expansion of NK cells would alleviate lupus-like disease features in the bm12-based cGvHD mouse model. Unexpectedly, both therapeutic and prophylactic administration of these cytokine-based therapeutics at least transiently enhanced disease measures. Through depletion studies, CD8 T cells were implicated in the disease aggravating activity of these cytokine-based therapeutics. These results suggest that activation and expansion of CD8 T cells may be unfavorable in the context of lupus, which is consistent with the correlation between elevated CD8 T-cell frequencies and poor disease prognosis in patients with lupus ^55^. Our findings represent pertinent discoveries regarding the dynamics between NK cells, CD8 T cells, and systemic autoimmune disease in mice.

Similar to the paucity of circulating NK cells in patients with lupus and reduced numbers of NK cells in other mouse models of lupus-like disease ^11-16,37,56,57^, we observe a marked contraction of the NK-cell compartment in mice subjected to the bm12 cGvHD model. Though the cause of this contraction remains undefined, we find that prophylactic treatment of diseased mice with either IL-2C or N-803 can counteract NK-cell loss and even promote expansion of the NK-cell pool. Coupled with experiments detailing IL-2-based therapeutics in the MRL/lpr and B6 x DBA mouse models ^28,29^, these data support the conclusion that therapeutic expansion of NK cells is possible in the context of lupus-like disease.

Unfortunately, application of IL-2C or N-803 to the bm12 cGvHD model of lupus at least transiently exacerbated disease. In both treatment approaches, the expansion of CD8 T cells was implicated in exaggerated bm12-engraftement-induced GC responses, as supported by CD8-depletion studies. Though the anti-CD8α antibody used to deplete CD8 T cells can also deplete CD8α^+^ dendritic cells ^58^, dendritic cells are not known to respond to N-803 ^53^. Dendritic cells are thus unlikely to be a confounding driver of N-803-exacerbated disease that can be reversed with CD8α-depleting antibody. However, as N-803 can promote secretion of IFN-γ from CD8 T cells ^31,39,59,60^, and IFN-γ can cause pathologic accumulation of T_FH_ cells and GCs ^61-63^, we speculate that excess IFN-γ production after N-803 or IL-2C therapy may provoke worse disease outcomes, a possibility that we will continue to explore. Regardless of the mechanism by which CD8 T cells may aggravate disease measures in this model, it is interesting to note that our results contrast with those obtained following therapeutic application of IL-2C to the B6 × DBA/2 F1 cGvHD mouse model of lupus-like disease ^29^, in which IL-2C-stimulated CD8 T cells ameliorated disease. This dichotomy is likely explained by the role of additional class I MHC mismatches in the B6 x DBA/2 F1 model that are not present in the bm12 model. In B6 x DBA/2 F1 system, IL-2C enhancement of CD8 T cells would lead to improved rejection of class I-mismatched cells, thereby curtailing the cGvHD response driving lupus-like disease. A similar elimination of pathogenic cells may drive the reduction in disease observed after IL-2C therapy of lupus-like disease in MRL/*lpr* mice.

While evidence for beneficial functions of NK cells in lupus remains scarce, our data suggest that specific expansion of NK cells (in the context of CD8 T cell depletion) is not necessarily detrimental to disease progression. The potential for increased clinical safety of NK cell-based versus T-cell based immunotherapy strategies has already been established in the context of chimeric antigen receptor cellular therapy of cancer ^64^. Since no mouse model of lupus-like disease accurately reflects all of the parameters driving human lupus, these studies collectively suggest that cytokine-mediated expansion of effector lymphocytes may still hold promise in the clinic if the dueling roles of NK cells and CD8 T cells in lupus-like disease can be better delineated. The advancement and further testing of NK-cell expansion for autoimmune disease will likely require development of reagents that specifically target NK cells and an improved understanding of the factors driving specific subsets of immunoregulatory NK cells.

Collectively, our data suggests that treatment of lupus with immunotherapeutics that selectively expand endogenous CD8 T and NK cells should be approached with caution and may not achieve the therapeutic effect of similar agents that has been observed in multiple sclerosis. Moreover, the increased risk of hematologic malignancy in patients with lupus ^65^ may complicate clinical application of N-803 or other antitumor therapeutics with the potential to aggravate autoimmune sequelae in these patients. On the other hand, NK cells are potent antitumor effector cells, and selective expansion of these cells without concurrent autoimmune-exacerbation by T cells may provide clinically valuable anti-leukemic effects even if these NK cells are unable to alleviate underlying elements of lupus-like disease.

## Acknowledgements

We would like to thank J. Klarquist and E. Janssen for expertise and for generating the Bm12/CD45.1 mouse line that was used for our disease model. We also thank Q. Ma, P. Devarajan, and W. Shao for assistance in evaluating lupus-like disease. Investigators on this project were supported by National Institutes of Health (NIH) grants DA038017, AI148080, AR073228, AI145304 (S.N.W.), T32GM063483, T32AI118697 (S.D.R. and A.A.), AR071651, and AR076316 (H.I.B.). Additional support provided to S.N.W. and H.I.B. by Cincinnati Children’s Research Foundation, the Cincinnati Center for Pediatric Genomics, and the Dr. Ralph and Marian Falk Medical Research Trust. S.D.R. was also supported by a Gina Finzi Memorial Student Fellowship from the Lupus Foundation. D.E.O. is supported by an American Heart Association Fellowship and the Arnold Strauss Fellowship. A.A. is supported by the Albert Ryan Fellowship. The Cincinnati Children’s Flow Cytometry Core is supported by NIH grants AR047363, AR070549, DK078392, DK090971, S10OD025045 and S10OD023410. N-803 and guidance in use of this drug were kindly provided by the team at ImmunityBio (formerly known as Altor Bioscience).

